# Distinct Tumor Necrosis Factor Alpha Receptors Dictate Stem Cell Fitness Versus Lineage Output in *Dnmt3a*-Mutant Clonal Hematopoiesis

**DOI:** 10.1101/2022.07.03.498502

**Authors:** Jennifer M. SanMiguel, Elizabeth Eudy, Matthew A. Loberg, Kira A. Young, Jayna J. Mistry, Logan S. Schwartz, Tim Stearns, Grant A. Challen, Jennifer J. Trowbridge

## Abstract

Clonal hematopoiesis resulting from enhanced fitness of mutant hematopoietic stem cells (HSCs) associates with both favorable and unfavorable health outcomes related to the types of mature mutant blood cells produced, but how this lineage output is regulated is unclear. Using a mouse model of a clonal hematopoiesis-associated mutation, *DNMT3A*^R882/+^ (*Dnmt3a*^R878H/+^), we found that aging-induced TNFα signaling promoted the selective advantage of mutant HSCs as well as stimulated mutant B lymphoid cell production. Genetic loss of TNFα receptor TNFR1 impaired mutant HSC fitness without altering lineage output, while loss of TNFR2 reduced lymphoid cell production and favored myeloid cell production from mutant HSCs without altering overall fitness. These results support a model where clone size and mature blood lineage production can be independently controlled to harness potential beneficial aspects of clonal hematopoiesis.

**Statement of Significance:** Through identification and dissection of TNFα signaling as a key driver of murine *Dnmt3a*-mutant hematopoiesis, we report the discovery that clone size and production of specific mature blood cell types can be independently regulated.

## Introduction

Clonal hematopoiesis is an aging-associated condition wherein hematopoietic stem cells (HSCs) have acquired a somatic mutation or copy number alteration that places them at a selective advantage. Both favorable and unfavorable health conditions have been associated with clonal hematopoiesis. Large clone size (VAF > 0.02) is associated with increased risk of hematologic malignancy, atherosclerosis, cardiovascular disease, type 2 diabetes, and osteoporosis^1–3^. Many of these conditions have been related to abnormal production of pro-inflammatory myeloid cell types such as macrophages and mast cells^4–7^. However, clonal hematopoiesis is naturally found in very aged populations without compromising survival^8^ and is associated with reduced risk of Alzheimer’s disease^9^. Furthermore, clonal hematopoiesis driven by mutations in the DNA methyltransferase *DNMT3A* has been associated with increased survival of recipients after bone marrow transplantation related to mutant T lymphoid cell production^10^ and maintenance of functional T cell immunity in a supercentenarian^11^. Thus, rather than developing methods to reduce clonal hematopoiesis altogether, further understanding of the molecular basis of both clone size and lineage potential will empower strategies to harness beneficial aspects of clonal hematopoiesis while reducing adverse health risks. Here, we used a mouse model of a clonal hematopoiesis-associated mutation in *DNMT3A* (*Dnmt3a*^R878H/+^)^12^ to study the molecular basis of HSC competition and lineage output.

## Results

Our group recently found that the middle-aged bone marrow (BM) microenvironment drives HSC aging^13^. This work established an experimental paradigm to evaluate potency of *Dnmt3a*^R878H/+^ HSCs in the aged BM microenvironment and identify HSC-extrinsic factors that modulate their selective advantage. We transplanted *Dnmt3a*^R878H/+^ HSCs into young and middle-aged recipient mice (Fig. 1A). *Dnmt3a*^R878H/+^ HSCs transplanted into aged recipients generated greater long-term multilineage hematopoiesis compared to control HSCs (Fig. 1B-C and Supplementary Fig. 1A) and gave rise to expanded HSC and multipotent progenitor (MPP) populations (Fig. 1D-E and Supplementary Fig. 1B). In addition, aged recipients had higher proportions of *Dnmt3a*^R878H/+^ megakaryocyte and erythroid-primed MPPs (MPP^Mk/E^) and lymphoid-primed MPPs (MPP^Ly^) (Fig. 1E). The latter was consistent with increased mature *Dnmt3a*^R878H/+^ B lymphoid cells (Supplementary Fig. 1A). No change in frequency of mature T lymphoid cells was observed, which may be explained in part by thymic involution during aging^14^. Control mice did not show significant MPP^Ly^ increase nor consistently higher mature B lymphoid cell production (Fig. 1E and Supplementary Fig. 1A-B). Together, these results demonstrate that the aged BM microenvironment provides a context in which *Dnmt3a*^R878H/+^ HSCs have enhanced selective advantage over wild-type HSCs as well as altered lineage output.

**Figure. 1.**
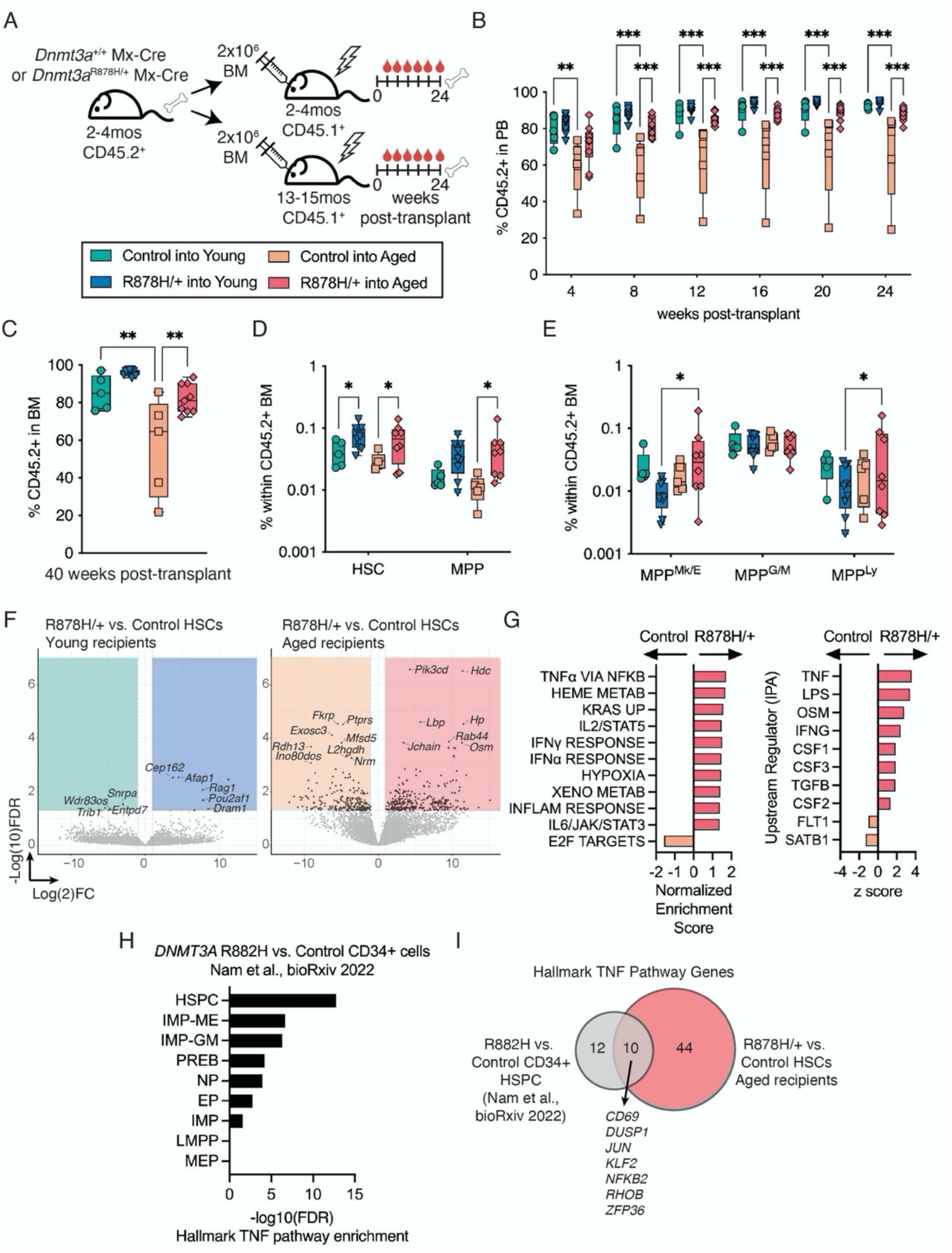
*Dnmt3a*^R878H/+^ HSCs Engage a TNFα-Induced Program in the Aged BM Microenvironment that is Conserved in Human *DNMT3A*-Mutant Clonal Hematopoiesis. (**A**) Schematic of experimental design to compare Mx-Cre control and *Dnmt3a*^R878H/+^ (R878H/+) engraftment in young (2-4mo) and aged (13-15mo) recipient mice. (**B**) Frequency of donor (CD45.2^+^) cells in peripheral blood (PB) of recipient mice post-transplant. Significance calculated using two-way ANOVA with Tukey’s multiple comparisons test. (**C**) Frequency of donor cells in bone marrow (BM) of recipient mice. Significance calculated using one-way ANOVA with Bonferroni’s multiple comparisons test. (**D**) Frequency of HSCs and MPPs in donor-derived BM cells. Significance calculated using two-way ANOVA with Fisher’s LSD. (**E**) Frequency of MPP^Mk/E^, MPP^G/M^ and MPP^Ly^ in donor-derived BM cells. Significance calculated using two-way ANOVA with Fisher’s LSD. (**F**) Volcano plots with significantly differentially expressed genes (FDR < 0.5, logFC > 2) within colored boxes (*n* = 2-4 biological replicates). (**G**) Enrichment of hallmark gene sets (left panel) and predicted upstream regulators (right panel) in control vs. *Dnmt3a*^R878H/+^ HSCs in aged recipient mice. (**H**) Hallmark TNF pathway enrichment across stem and progenitor populations between human *DNMT3A*R882H vs control CD34+ cells. (**I**) Overlap of differentially expressed genes in *DNMT3A*R882H vs control CD34+ cells and R878H/+ vs aged mouse HSCs. (**B-E**) Dots represent individual recipient mice, boxes show 25 to 75^th^ percentile, line is median, whiskers show min to max. \**P* < 0.05, *\*\*P* < 0.01, *\*\*\*P* < 0.001.

To identify molecular signatures underlying expanded *Dnmt3a*^R878H/+^ hematopoiesis in the aged BM microenvironment, we performed RNA-seq on independent biological replicates of HSCs re-isolated from young and aged recipient mice. Our experimental design specified only a sublethal dose of irradiation to recipient mice, to better preserve HSC-extrinsic signals from the BM microenvironment^15–19^ (Supplementary Fig. 1C). A greater number of differentially expressed genes in *Dnmt3a*^R878H/+^ vs. control HSCs were found in aged compared to young recipient mice (Fig. 1F). Using gene and pathway enrichment analyses, TNFα was identified as the top enriched gene signature and predicted upstream regulator in *Dnmt3a*^R878H/+^ HSCs in aged mice (Fig. 1G). A previously described TNFα-induced transcriptional program in HSCs, enriched in pro-survival genes^20^, was elicited in *Dnmt3a*^R878H/+^ HSCs in aged mice (Supplementary Fig. 1D). In addition, *Dnmt3a*^R878H/+^ HSCs in aged mice maintained expression of transcriptional programs that define mouse and human HSCs^21,22^ (Supplementary Fig. 1D). Consistent with these observations, human *DNMT3A*^R882H/+^ CD34^+^ HSPCs showed enrichment of a TNFα pathway signature compared to *DNMT3A*^+/+^ CD34^+^ HSPCs isolated from the same individuals (Fig. 1H)^23^. Several TNFα target genes were commonly upregulated in human *DNMT3A*^R882H/+^ CD34^+^ HSPCs and mouse *Dnmt3a*^R878H/+^ HSCs, including *JUN* and *NFKB2* (Fig. 1I). Taken together, the selective advantage of mouse and human *DNMT3A*-mutant HSCs correlates with a TNFα-induced, HSC-survival-associated gene expression signature.

To assess the extent to which TNFα directly promotes young *Dnmt3a*-mutant HSC survival, we added recombinant TNFα to mixed cultures of wild-type and *Dnmt3a*-mutant HSCs in media that sustains HSC self-renewal^24,25^ (Fig. 2A). TNFα treatment reduced the number of control but not *Dnmt3a*^R878H/+^ cells produced over the culture period (Supplementary Fig. 2A) and did not alter stem/progenitor cell surface marker phenotypes (Supplementary Fig. 2B). Post culture, cells were transplanted into recipient mice to assess HSC function. TNFα-treated control HSCs did not sustain long-term multilineage engraftment (Fig. 2B and Supplementary Fig. 2C-F). In contrast, TNFα-treated *Dnmt3a*^R878H/+^ HSCs increased production of mature hematopoietic cells in the short term (4 weeks post-transplant) followed by sustained multilineage engraftment at levels comparable to vehicle-treated *Dnmt3a*^R878H/+^ cells. In addition, TNFα stimulation transiently increased *Dnmt3a*^R878H/+^ B lymphoid cell production (Fig. 2C-D and Supplementary Fig. 2C), in contrast to transient myeloid regeneration from TNFα-treated control HSCs as has been previously reported^20^. In the bone marrow of mice transplanted with TNFα-treated control HSCs, we observed trends toward reduced HSC, MPP^Mk/E^, and MPP^G/M^ populations (Supplementary Fig. 2G-H). This decrease was not observed in TNFα-treated *Dnmt3a*^R878H/+^ HSCs. Thus, the ability of TNFα to promote myeloid regeneration at the expense of maintaining self-renewal in control HSCs is disrupted in *Dnmt3a*^R878H/+^ HSCs. Instead, TNFα-treated *Dnmt3a*^R878H/+^ HSCs favor lymphoid regeneration and maintain their self-renewal. To assess if our observations were specific to using *Mx*-Cre recombination and/or the *Dnmt3a*^R878H^ mutation, we performed TNFα stimulation of HSCs from tamoxifen-inducible *Fgd5*-Cre-driven *Dnmt3a*^R878H/+^ mice as well as germline *Dnmt3a*^+/−^ mice (Fig. 2E). We find that TNF treatment reduced the number of control cells but not *Fgd5*-Cre-driven *Dnmt3a*^R878H/+^ cells (Fig. 2F) or *Dnmt3a*^+/−^ cells (Fig. 2G) over the culture period. After transplant into recipient mice, TNFα-treated *Dnmt3a*^+/−^ HSCs vs. control HSCs had increased production of mature hematopoietic cells in the short term (Fig. 2H) and increased B lymphoid relative to myeloid cell production (Fig. 2I). Thus, disrupted myeloid regeneration and HSC survival phenotypes induced by TNFα are broadly relevant to *Dnmt3a*-mutant clonal hematopoiesis.

**Figure 2.**
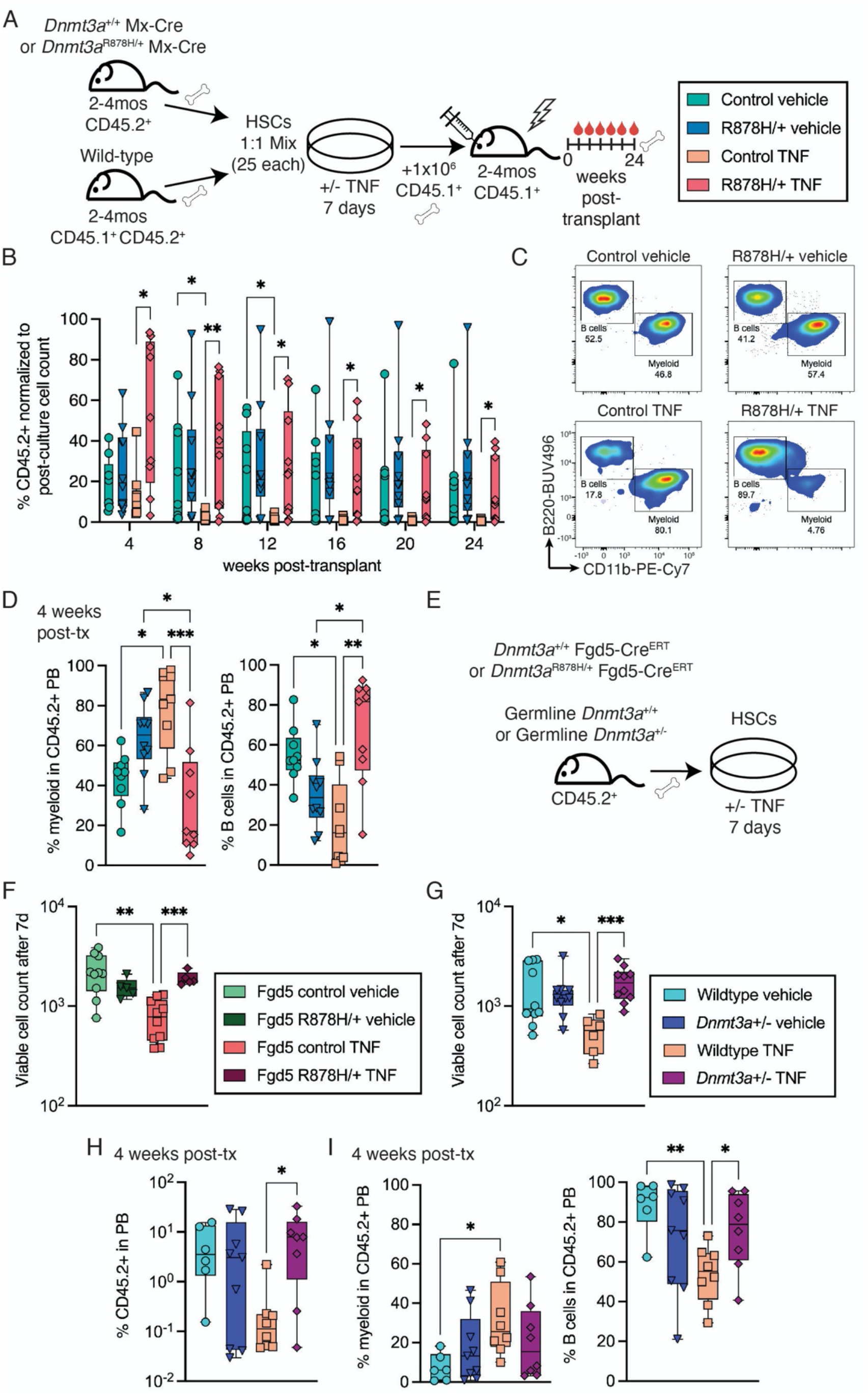
*Dnmt3a*-Mutant HSCs Maintain Self-Renewal and Generate B Lymphoid Cells Following TNF Stimulation. (**A**) Schematic of experimental design to test response of Mx-Cre control and *Dnmt3a*^R878H/+^ (R878H/+) HSCs to recombinant TNFα *ex vivo* under growth conditions that favor HSC expansion. (**B**) Normalized frequency of donor-derived cells in PB of recipient mice posttransplant. Significance calculated using mixed-effects model with Fisher’s LSD. (**C**) Representative flow cytometry plots showing B cell and myeloid cell frequencies in donor derived PB at 4 weeks posttransplant. (**D**) Frequency of myeloid (left) and B cells (right) in donor derived PB at 4 weeks posttransplant. Significance calculated using two-way ANOVA with Sidak’s multiple comparison’s test (**E**) Schematic of experimental design to test TNFα response of Fgd5-Cre^ERT^ control vs. Fgd5-Cre^ERT^ *Dnmt3a*^R878H/+^ (R878H/+) HSCs, and germline *Dnmt3a*^+/+^ vs. *Dnmt3a*^+/−^ HSCs. (**F**, **G**) Viable cell counts after 7 days of culture. Significance calculated using Brown-Forsythe and Welch ANOVA with Welch’s correction. (**H**) Frequency of donor-derived cells in PB of recipient mice at 4 weeks post-transplant. Significance calculated using one-way ANOVA with Fisher’s LSD. (**I**) Frequency of myeloid (left) and B cells (right) in donor derived PB at 4 weeks post-transplant. Significance calculated using one-way ANOVA with Fisher’s LSD. (**B**, **D**, **F-I**) Dots represent individual recipient mice, boxes show 25 to 75^th^ percentile, line is median, whiskers show min to max whiskers show min to max. \**P* < 0.05, \*\**P* < 0.01, \*\*\**P* < 0.001.

TNFα signaling occurs through two distinct TNFα receptors, TNFR1 (*Tnfrsf1a*) and TNFR2 (*Tnfrsf1b*). Both TNFR1 and TNFR2 are expressed on HSC and MPP populations and are not altered in expression in *Dnmt3a*^R878H/+^ mice (Supplementary Fig. 3A-D). To determine which of these receptors are responsible for TNFα-mediated selective advantage of *Dnmt3a*^R878H/+^ HSCs and B lymphoid cell production, we crossed *Dnmt3a*^R878H/+^ mice with *Tnfrsf1a* or *Tnfrsf1b* knockout mice (Supplementary Fig. 3E)^26^ and rigorously tested HSC function using competitive serial BM transplantation into aged recipients (Fig. 3A). Loss of TNFR1, but not TNFR2, eliminated the selective advantage of *Dnmt3a*^R878H/+^ PB and BM cells in primary (Supplementary Fig. 4A-B) and secondary (Fig. 3B and Supplementary Fig. 5A) transplant, including in the HSC compartment itself (Supplementary Fig. 4E-F and Supplementary Fig. 5E-F). No change in engraftment was observed in TNFR1 knockout-only controls (Supplementary Fig. 6A-B and Supplementary Fig. 7A, E-G), demonstrating that this is a specific dependency of *Dnmt3a*^R878H/+^ cells. In contrast, loss of TNFR2, but not TNFR1, reduced the proportion of B and T lymphoid cells and increased the proportion of myeloid cells (Fig. 3C-D and Supplementary Fig. 4C-D and Supplementary Fig. 5B-D) without altering overall white blood cell production (Supplementary Fig. 5B). This myeloid-biased hematopoiesis was also observed in TNFR2 knockout-only controls but only late in the post-secondary-transplant period (Supplementary Fig. 6C-D and Supplementary Fig. 7C-D) and was more mild as it did not increase neutrophil count (Supplementary Fig. 7B), demonstrating that TNFR2 signaling regulates lymphoid cell output from *Dnmt3a*^R878H/+^ cells more so than wild-type cells. Taken together, our results demonstrate that distinct TNFα receptor signaling controls *Dnmt3a*^R878H/+^ HSC self-renewal and regenerative capacity versus lineage output in response to elevated TNFα. To test if pharmacological blockade of TNF signaling would recapitulate TNFR knockout phenotypes, we treated competitive secondary transplant mice with etanercept, a pan-TNFα inhibitor (Fig. 3 E). Etanercept treatment reduced competitive engraftment of *Dnmt3a*^R878H/+^ cells in the PB (Fig. 3F) and trended toward reduction in the BM (Fig. 3G), and reduced the frequency of *Dnmt3a^R878H/+^* HSC, MPP^Mk/E^, and MPP^Ly^ populations at the time of harvest (Fig. 3H). In contrast, *Dnmt3a*^R878H/+^ myeloid-primed multipotent progenitors MPP^G/M^ trended toward increase after etanercept treatment. These results suggest that pan-TNF inhibition results in a mix of our observed TNFR knockout phenotypes, that is, reduced selective advantage of *Dnmt3a*^R878H/+^ hematopoiesis as well as myeloid lineage bias at the stem/progenitor cell level.

**Figure 3.**
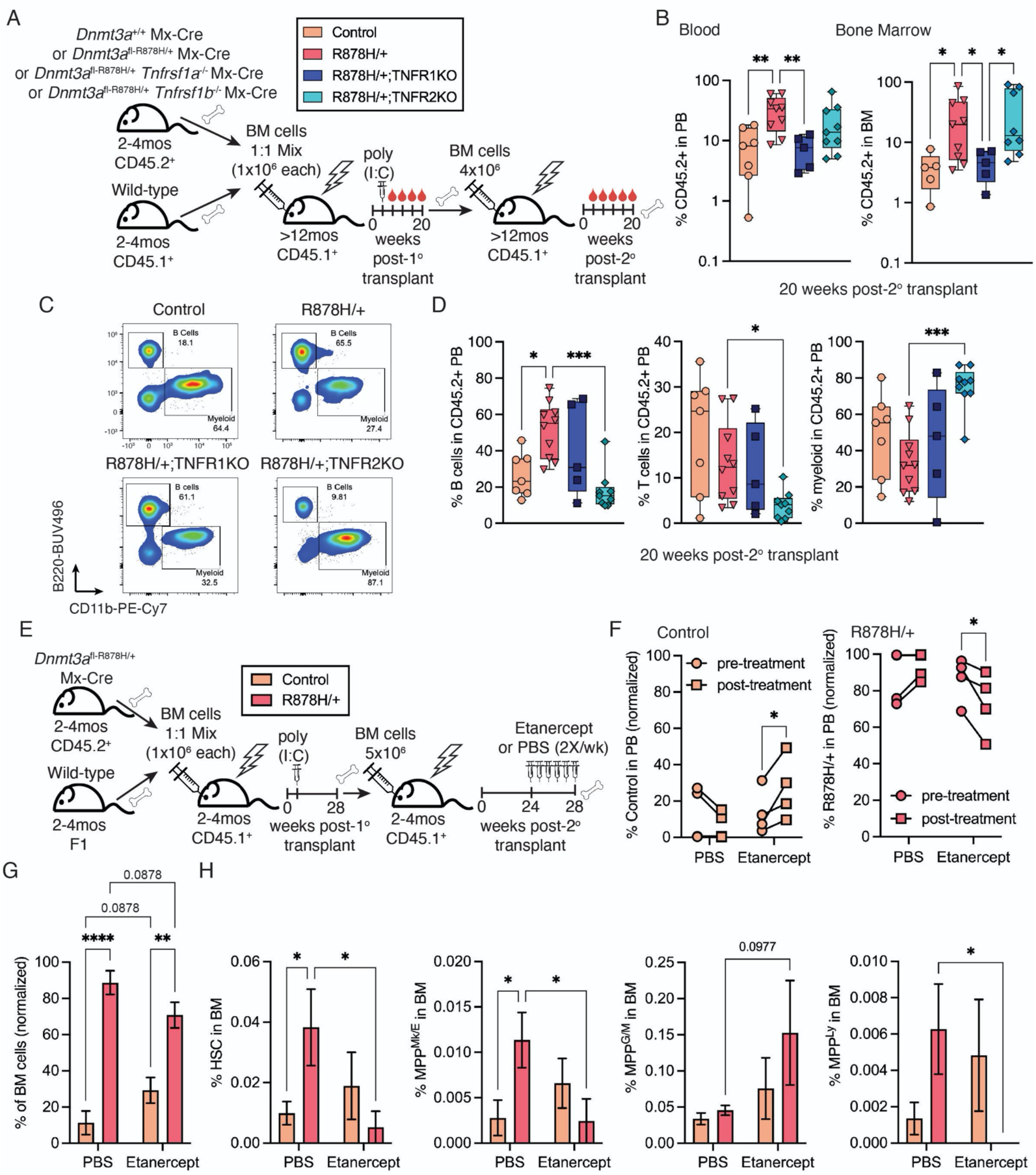
TNFR1 is Required for *Dnmt3a*^R878H/+^ HSC Self-Renewal While TNFR2 Regulates Lymphoid Cell Production. (**A**) Schematic of experimental design to test competitive, serial transplant of Mx-Cre control, *Dnmt3a*^R878H/+^ (R878H/+), *Dnmt3a*^R878H/+^ *Tnfrsf1a*^-/-^ (R878H/+;TNFR1KO) and *Dnmt3a*^R878H/+^ *Tnfrsf1b*^-/-^ (R878H/+;TNFR2KO) in aged (>12mo) recipient mice. (**B**) Frequency of donor cells in PB (left) and BM (right) of recipient mice at 20 weeks post-secondary transplant. Significance was calculated using Brown-Forsythe and Welch ANOVA with Welch’s correction. (**C**) Representative flow cytometry plots showing B cell and myeloid cell frequencies in donor derived PB. (**D**) Frequency of B cells (left), T cells (center) and myeloid cells (right) in donor derived PB at 20 weeks post-secondary transplant. Significance was calculated using one-way ANOVA with Tukey’s multiple comparisons test. (**E**) Experimental schematic of transplant experiment with etanercept treatment. (**F**) Frequency of control or R878H/+ donor cells in PB pre-and post-etanercept treatment. Significance was calculated using two-way ANOVA with Fisher’s LSD. (**G**) Frequency of control or R878H/+ donor cells in BM post-etanercept or vehicle treatment. Significance was calculated using two-way ANOVA with Fisher’s LSD. (**H**) Frequency of HSC, MPP^Mk/E^, MPP^G/M^, and MPP^Ly^ populations in control or R878H/+ BM. Significance was calculated using two-way ANOVA with Fisher’s LSD. (**B, D**) Dots represent individual recipient mice, boxes show 25 to 75^th^ percentile, line is median, whiskers show min to max. (**F**) Dots represent mean across biological replicates (*n* = 4). (**G, H**) Bars represent mean +/-SEM (*n* = 4). \**P* < 0.05, \*\**P* < 0.01, \*\*\**P* < 0.001, \*\*\*\**P* < 0.0001.

To interrogate mechanisms by which TNFα signaling through different receptors impacts *Dnmt3a*^R878H/+^ cells, we harvested donor-derived hematopoietic stem and progenitor cells from secondary transplant recipient mice for single cell RNA-sequencing (*n* = 3-4 biological replicates per genotype) (Fig. 4A). After quality control filtering (Supplementary Fig. 8A-C), a total of 64,830 cells clustered into 22 populations (Fig. 4B). These clusters were classified based on published data to identify HSC, multipotent progenitor, and lineage-specified progenitor^27^ (Supplementary Fig. 8D and Supplementary Table 1). Pseudotime trajectory analysis revealed that myeloid progenitor differentiation from HSCs (My) followed a distinct path in *Dnmt3a*^R878H/+^ vs. control BM (Fig. 4C), whereas erythroid (Ery) and megakaryocyte (Mk) progenitor differentiation paths were similar. Loss of TNFR1 fully corrected the aberrant *Dnmt3a*^R878H/+^ HSC to myeloid progenitor differentiation trajectory to closely resemble control BM, while loss of TNFR2 created multiple myeloid differentiation trajectories from *Dnmt3a^R878H/+^* HSCs. These data are highly consistent with our functional observations that loss of TNFR1 eliminates *Dnmt3a*^R878H/+^ selective advantage and loss of TNFR2 biases toward myeloid cell production. Within the subsets of hematopoietic stem and progenitor cells that we identified, TNF signaling was most strongly enriched in *Dnmt3a*^R878H/+^ vs. control HSCs (Fig. 4D), supporting that TNF-induced phenotypes are initiated at the HSC level. Indeed, many downstream TNF targets were increased in expression in *Dnmt3a*^R878H/+^ vs. control HSCs, and several of these target genes are known to be hypomethylated in *Dnmt3a*^R878H/+^ HSCs^28^ (Fig. 4E). *Dnmt3a*^R878H/+^ TNFR1 knockout HSCs and *Dnmt3a*^R878H/+^ TNFR2 knockout HSCs demonstrated downregulation of *Tnfrsf1a* and *Tnfrsf1b* transcripts, respectively.

**Figure 4.**
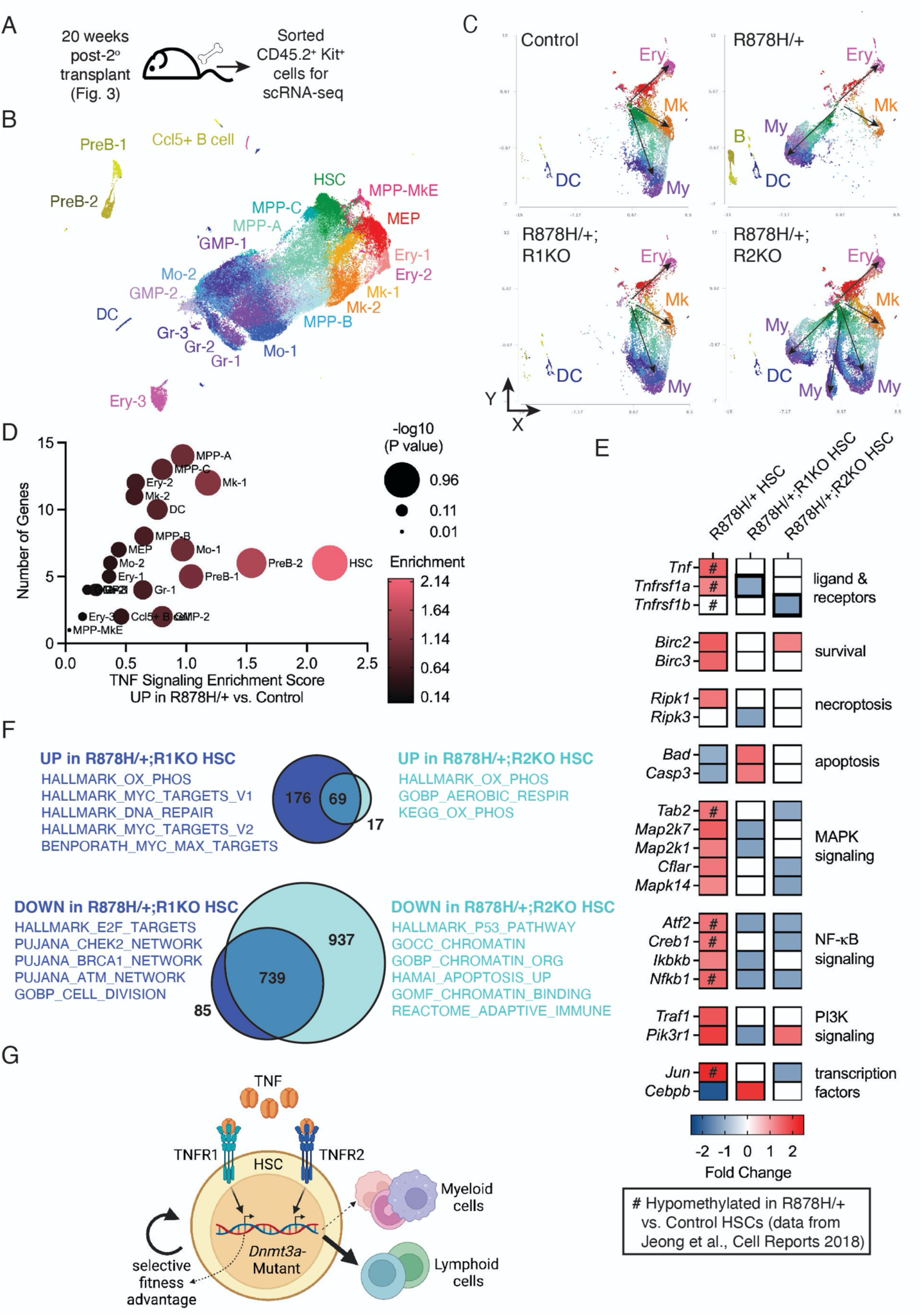
TNFR1 and TNFR2 Engage Distinct Transcriptional Programs in *Dnmt3a*^R878H/+^ HSCs. (**A**) Experimental schematic for single cell (sc) RNA-seq. Data was collected from independent biological replicates of Mx-Cre control (*n* = 3), *Dnmt3a*^R878H/+^ (R878H/+) (*n* = 4), *Dnmt3a*^R878H/+^ *Tnfrsf1a*^-/-^ (R878H/+;R1KO) (*n* = 3) and *Dnmt3a*^R878H/+^ *Tnfrsf1b*^-/-^ (R878H/+;R2KO) (*n* = 4). (**B**) UMAP projection of combined data identifying 22 cell clusters. (**C**) Pseudotime visualization showing predicted differentiation trajectories from HSCs to erythroid (Ery), megakaryocyte (Mk), myeloid (My), B cell (B) and dendritic cell (DC) lineages in each genotype pool. (**D**) TNF signaling enrichment score in R878H/+ vs. control cell clusters. (**E**) Heatmap representing fold change in expression of *Tnf, Tnfrsf1a* (TNFR1), *Tnfrsf1b* (TNFR2), and downstream TNF-regulated genes comparing R878H/+ vs. control HSCs, R878H/+;R1KO vs. R878H/+ HSCs, and R878H/+;R2KO vs. R878H/+ HSCs. “#” indicate loci hypomethylated in R878H/+ vs. control HSCs^28^. (**F**) Venn diagrams of overlap between upregulated genes (top) and downregulated genes (bottom) in R878H/+;R1KO and R878H/+;R2KO HSCs compared to R878H/+ HSCs. From each comparison, unique gene lists were used to determine gene signature enrichment. (**G**) Working model created with BioRender.com. TNFα-TNFR1 signaling dictates *Dnmt3a-mutant* HSC self-renewal whereas TNFα-TNFR2 signaling promotes lymphoid lineage cell production. Decline in TNFα-TNFR2 signaling results in unrestrained production of *Dnmt3a*-mutant myeloid cells.

Focusing on unique transcriptional changes in *Dnmt3a*^R878H/+^ TNFR1 vs. TNFR2 knockout HSCs, we found that loss of TNFR1 resulted in increased expression of mediators of apoptosis, initiation factors for DNA repair and checkpoint activation, increased expression of *Cebpb*, and decreased cell division (Fig. 4E-F). In contrast, loss of TNFR2 resulted in increased expression of the apoptosis inhibitor *Birc2*, decrease in tumor suppressor p53, dysregulation of chromatin organization and decreased B and T lymphoid ‘adaptive immune’ signatures. Together, our work supports that TNFα-TNFR1 signaling promotes *Dnmt3a*^R878H/+^ HSC competitive advantage through evasion of apoptosis, accumulation of DNA damage, self-renewal, and cell cycling. In contrast, TNFα-TNFR2 signaling promotes lymphoid cell production from *Dnmt3a*^R878H/+^ HSCs, and restrains myeloid cell production, through chromatin regulation and expression of lymphoid-specifying genes.

## Discussion

Inhibition of pro-inflammatory cytokines, including TNFa, has been proposed as a generalizable strategy to reduce fitness of CH-mutant HSCs and risk of CH-associated disease states as these are related to abnormal production of pro-inflammatory myeloid cell types^7,29–32^. Our work suggests that pan-TNF inhibition can reduce *Dnmt3a*^R878H/+^ HSC fitness but also results in more complex and potentially detrimental effects due to unrestrained *Dnmt3a*^R878H/+^ myeloid cell production. Alternatively, we have found that programs dictating *Dnmt3a*-mutant HSC fitness and mature hematopoietic lineage cell production can be separated by downstream pathway dissection of TNFa signaling. Our work supports the possibility of independently regulating clone fitness and lineage output to reduce risk of CH-associated diseases such as hematologic malignancy and harness potential beneficial aspects of CH such as maintained T-cell function in aging, survival of stem cell transplant recipients, and reduced risk of Alzheimer’s disease.

## Materials and Methods

### Experimental Animals

C57BL/6J (stock #00664, referred to as “CD45.2^+^”) and B6.SJL-Ptprca Pepcb /BoyJ (stock #002014, referred to as “CD45.1^+^”) mice were obtained from, and aged within, The Jackson Laboratory (JAX). *Dnmt3a*^fl-R878H/+^ mice^12^ were crossed to B6.CgTg(Mx1-cre)1Cgn/J mice (referred to as Mx-Cre)^33^. *B6.129S-Tnfrsf1b^tm1/mx^*; *Tnfrsf1a^tm1/mx^* ^26^ were obtained from JAX (stock #003243) and crossed to *Dnmt3a* ^fl-R878H/+^;Mx-Cre. The Jackson Laboratory’s Institutional Animal Care and Use Committee (IACUC) approved all experiments.

### Flow Cytometry and Cell Sorting

Single cell suspensions of BM were prepared by filtering crushed, pooled femurs, tibiae, and iliac crests from each mouse. BM mononuclear cells (MNCs) were isolated by Ficoll-Paque (GE Healthcare Life Sciences) density centrifugation and stained with a combination of fluorochrome-conjugated antibodies from eBioscience, BD Biosciences, or BioLegend: CD45.1 (clone A20), CD45.2 (clone 104), c-Kit (clone 2B8), Sca-1 (clone 108129), CD150 (clone TC15-12F12.2), CD48 (clone HM48-1), FLT3 (Clone A2F10), CD34 (clone RAM34), FcgR (clone 2.4G2), mature lineage (Lin) marker mix and a viability stain. Stained cells were sorted using a FACSAria or a FACSymphony S6 Sorter (BD Biosciences), or analyzed on a FACSymphony A5 or LSR II with Diva software (BD Biosciences) based on the following surface marker profiles: HSC (Lin-Sca-1+ c-Kit+ Flt3-CD150+ CD48-), MPP (Lin-Sca-1+ c-Kit+ Flt3-CD150-CD48-), MPP^Mk/E^ (Lin-Sca-1+ c-Kit+ Flt3-CD150+ CD48+), MPP^G/M^ (Lin-Sca-1+ c-Kit+ Flt3-CD150-CD48+), MPP^Ly^ (Lin-Sca-1+ c-Kit+ Flt3+), HSC+MPP (Lin-Sca-1+ c-Kit+), and MyPro (Lin-Sca-1-c-Kit+). TNFR1 staining used a tertiary staining method using purified anti-mouse TNFR1 (Biolegend cat. #113001) followed by biotin goat-anti-hamster IgG (Biolegend cat #405501) followed by streptavidin-PE (BD cat. #554061). TNFR2 staining used a secondary staining approach using anti-mouse TNFR2 (Biolegend cat no. 113403) followed by streptavidin-PE. PB samples were stained and analyzed using a cocktail of CD45.1, CD45.2, CD11b (clone M1/70), B220 (clone RA3-6B2), CD3e (clone 145-2C11), Ly6g (clone 1A8), and Ly6c (clone HK1.4) on an LSRII (BD) based on the following surface marker profiles: B cells (B220+ CD11b-CD3e-), T cells (CD3e+ B220-CD11b-), myeloid cells (CD11b+ B220-CD3-), granulocytes (CD11b+ B220-CD3-Ly6g+ Ly6c+), monocytes (CD11b+ B220-CD3-Ly6g-Ly6c+), macrophages (CD11b+ B220-CD3-Ly6g-Ly6c-). Gating analysis was performed using FlowJo software v10.

### Transplants into Young and Aged Recipient Mice

2–4-month-old *Dnmt3a*^+/+^ Mx-Cre or *Dnmt3a*^fl-R878H/+^ Mx-Cre donors were injected with poly(I:C). 2 months post-poly(I:C), 2 × 10^6^ post-ficoll whole bone marrow cells from the donors were transplanted into lethally irradiated (10 Gy) young (2-4 mos) or middle-aged (13-15 mos) CD45.1^+^ recipient mice. All transplant recipient mice were monitored every 4 weeks post-transplant by flow cytometry analysis of PB and were harvested for bone marrow analysis at 40 weeks post-transplant.

### Polyvinyl Alcohol (PVA) Culture and Transplants

2-month-old *Dnmt3a*^+/+^ Mx-Cre or *Dnmt3a*^fl-R878H/+^ Mx-Cre donors (CD45.2+) were injected with 5mg/kg poly(I:C) every other day for a total of five injections. 2 months post-poly(I:C), 25 CD45.2^+^ and 25 CD45.1^+^/CD45.2^+^ HSCs were sorted into a 96-well plate with Ham’s F12 media containing final concentrations of 1× Penicillin–streptomycin–glutamine (Gibco cat. # 10378-016), 10 mM HEPES (Gibco cat. #15630080), 1× Insulin–transferrin–selenium–ethanolamine (Gibco cat. #51500-056), 100 ng/mL recombinant murine TPO (Biolegend cat. # 593302), 10 ng/mL recombinant murine SCF (StemCell Technologies cat. # 78064), and 1 mg/mL polyvinyl alcohol (Sigma cat. # P8136)^24,25^ +/− 10 ng/mL recombinant murine TNF-α (PeproTech cat. #315-01A) and cultured for seven days at 37°C and 5% CO_2_. TNF-α was spiked into the cultures on day 4 and 6. On day 7, half of the wells were stained and analyzed by flow cytometry and half of the wells were harvested, mixed with 1 × 10^6^ CD45.1^+^ post-ficoll whole BM cells, and transplanted into young, lethally irradiated CD45.1^+^ recipients. PB was tracked monthly for 6 months post-transplant via flow cytometry.

### Etanercept Transplants

1×10^6^ BM cells from 2-4-month-old *Dnmt13a* Mx-Cre or *Dnmt3a*^fl-R878H/+^ MxCre donors were competitively transplanted with wild-type CD45.1+ CD45.2+ F1 BM cells in 2–4 month old CD45.1+ lethally irradiated recipients. Recipients were allowed to recover for one month and then poly(I:C) was administered every other day for a total of five injections to induce Cre expression. 28 weeks post-poly(I:C), bone marrow was harvested and 5 × 10^6^ whole bone marrow cells were transplanted into 2–4-month-old lethally irradiated CD45.1+ recipients. 24 weeks post-secondary transplant, etanercept (25 mg/kg, Millipore Sigma #Y0001969) or PBS was administered via IP twice per week for four weeks. PB was monitored weekly and at 28 weeks post-transplant, bone marrow was harvested for analysis.

### TNFR Knockout Transplants

1×10^6^ CD45.2+ cells were competed against 1×10^6^ CD45.1+ whole BM cells and transplanted into aged, lethally irradiated CD45.1+ recipient animals. One-month post-transplant, recipients received one IP injection of poly(I:C) and recombination was checked via PCR on PB. One month post poly(I:C), animals were bled monthly for 16 weeks. Bone marrow was harvested and 4×10^6^ whole BM cells were used for secondary transplantation into aged, lethally irradiated recipients. PB was analyzed starting at one-month post-transplant and continued monthly for 20 weeks. BM was harvested and analyzed by flow cytometry and Lin-c-kit+ CD45.2+ cells were FACS-sorted for single-cell RNA-sequencing. Complete blood counts (CBC) were performed on a Advia 120 Hematology Analyzer (Siemens).

### Bulk RNA-Sequencing and Analysis

2–4-month-old *Dnmt3a*^+/+^ Mx-Cre or *Dnmt3a*^R878H/+^ Mx-Cre donors were injected with poly(I:C) five times every other day. 1-month post-poly(I:C), 1×10^6^ whole BM cells from were transplanted into sublethally irradiated (6 Gy) young (2mos) or middle-aged (13-15 mos) CD45.1+ recipient mice. Recipients were harvested at 4 mos post-transplant for PB and BM analysis. CD45.2+ HSCs were sorted directly into RLT buffer (Qiagen) and flash frozen. Total RNA was isolated using the RNAeasy Micro Kit (Qiagen) including DNase treatment, and sample quality was assessed using a Nanodrop 2000 spectrophotometer (Thermo Scientific) and RNA 6000 Pico LabChip assay (Agilent Technologies). Libraries were prepared using the Ovation RNA-seq System V2 (NuGen) and Hyper Prep Kit (Kapa Biosystems). Library quality and concentration evaluated using D5000 ScreenTape assay (Agilent) and quantitative PCR (Kapa Biosystems). Libraries were pooled and sequenced 75bp single end on the NextSeq (Illumina) using NextSeq High Output Kit v2 reagents at a sequencing depth of >30 million reads per sample. Trimmed alignment files were processed using RSEM (v1.2.12). Alignment was completed using Bowtie 2 (v2.2.0). Expected read counts per gene produced by RSEM were rounded to integer values, filtered to include only genes that have at least two samples within a sample group having a cpm > 1, and were passed to edgeR (v3.14.0) for differential expression analysis. The GLM likelihood ratio test was used for differential expression in pairwise comparisons between sample groups which produced exact p-values per test. The Benjamini and Hochberg’s algorithm (p-value adjustment) was used to control the false discovery rate (FDR). Features with FDR-adjusted p-value < 0.05 were declared significantly differentially expressed. Differentially expressed genes were investigated for overlap with published datasets using Gene Set Enrichment Analysis (GSEA) and upstream regulators were predicted using Ingenuity Pathway Analysis (IPA) software (Qiagen).

### Single Cell RNA Sequencing and Analysis

Cells were counted on a Countess II automated cell counter (ThermoFisher) and 12,000 cells were loaded on to one lane of a 10X Chromium microfluidic chip. (10X Genomics). Single cell capture, barcoding and library preparation were performed using the 10X Chromium version 3.1 chemistry, according to the manufacturer’s protocol (#CG000315). cDNA and libraries were checked for quality on Agilent 4200 Tapestation and quantified by KAPA qPCR before sequencing; each gene expression library was sequenced at 18.75% of an Illumina NovaSeq 6000 S4 flow cell lane, targeting 6,000 barcoded cells with an average sequencing depth of 75,000 reads per cell. Illumina base call files for all libraries were demultiplexed and converted to FASTQs using bcl2fastq v2.20.0.422 (Illumina). The Cellranger pipeline (10× Genomics, version 6.0.0) was used to align reads to the mouse reference GRCm38.p93 (mm10 10× Genomics reference 2020-A), de-duplicate reads, call cells, and generate cell by gene digital counts matrices for each library. The resultant counts matrices were uploaded into PartekFlow (version 10.0.22.0428) for downstream analysis and visualization. This included log transformation of count data, principal component analysis, graph-based clustering from the top 20 principal components using the Louvain Algorithm, UMAP visualization, and pathway enrichment analysis. Trajectory and pseudotime analysis were performed using Monocle 3.

### Statistical Analysis

No sample group randomization or blinding was performed. All statistical tests, including evaluation of normal distribution of data and examination of variance between groups being statistically compared, were assessed using Prism 9 software (GraphPad).

## Data Availability Statement

All data in this study are deposited in the NCBI Gene Expression Omnibus (GEO) under accession number GSE189406 (bulk RNA-seq) and GSE203550 (single-cell RNA-seq).

## Acknowledgements

We thank all members of the Trowbridge lab for their help with experimental support and manuscript editing. We thank Ross Levine, Coleman Lindsley, and Michael Rauh for discussions and input, and Dan Landau and Neville Dusaj for providing single cell expression data from human *DNMT3A*-mutant CH samples. We thank the Scientific Research Services at The Jackson Laboratory, specifically the Single Cell Biology service, Histopathology-Clinical Chemistry, Genome Technologies, Computational Sciences, and Flow Cytometry. These shared services are supported in part by the JAX Cancer Center (P30 CA034196).

## Notes

This work was supported by NIH U01AG077925 (J.J.T.), NIH R01DK118072 (J.J.T.), NIH R01AG069010 (J.J.T.), and an EvansMDS Discovery Research grant (J.J.T.). This study was supported in part by The Jackson Laboratory’s Cancer Center Support Grant NIH P30 CA034196. J.M.S. was supported by NIH T32AG062409, NIH T32HD007065, and an American Society of Hematology (ASH) Scholar award. G.A.C. and J.J.T. are scholars of the Leukemia and Lymphoma Society. G.A.C. was supported by NIH R01 DK124883.

Conflict of interest disclosure: J.J.T. holds a sponsored research project with H3 Biomedicine. All other authors declare that they have no competing interests.

### Competing Interest Statement

J.J.T. holds a sponsored research project with H3 Biomedicine.

