## Supplemental Figures for "Distinct Tumor Necrosis Factor Alpha Receptors Dictate Stem Cell Fitness Versus Lineage Output in *Dnmt3a*-Mutant Clonal Hematopoiesis"

**SUPPLEMENTARY MATERIAL**

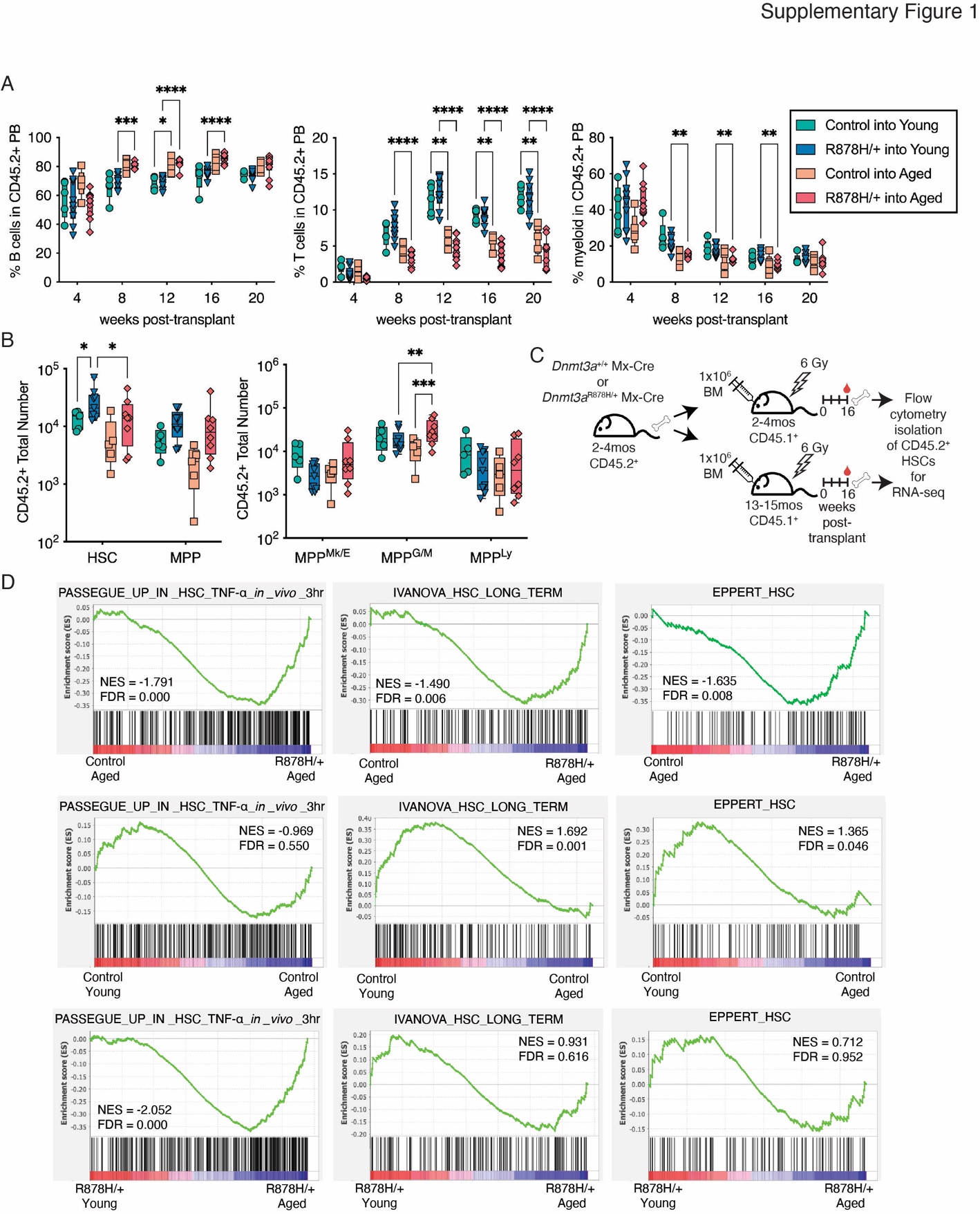

**Supplementary Figure 1. Lineage Composition and Gene Signature Enrichment in Control vs. *Dnmt3a*^R878H/+^ Hematopoiesis in Young and Aged Recipient Mice.** (**A**) Frequency of B cell (left), T cells (center) and myeloid cells (right) in donor derived PB. Significance was calculated using two-way ANOVA with Tukey’s multiple comparisons test. (**B**) Total number of CD45.2+ HSC and MPP (left) and MPP^Mk/E^, MPP^G/M^, MPP^Ly^ (right) in BM of recipient mice. Significance was calculated using two-way ANOVA with Fisher’s LSD. (**C**) Schematic of experimental design to re-isolate Mx-Cre control and *Dnmt3a*^R878H/+^ (R878H/+) HSCs from young and aged recipient mice for RNA-seq (*n* = 2-4 biological replicates per condition). (**D**) Enrichment of gene signatures in control vs. R878H/+ HSCs in aged recipient mice (top row), control HSCs in young vs. aged recipient mice (middle row), and R878H/+ HSCs in young vs. aged recipient mice (bottom row). These gene signatures define *in vivo* TNFα-stimulated target genes (left column), murine HSCs (center column), and human HSCs (right column). (**A, B**) Dots represent individual recipient mice, boxes show 25 to 75^th^ percentile, line is median, whiskers show min to max. **P* < 0.05, ***P* < 0.01, ****P* < 0.001, *****P* < 0.0001.

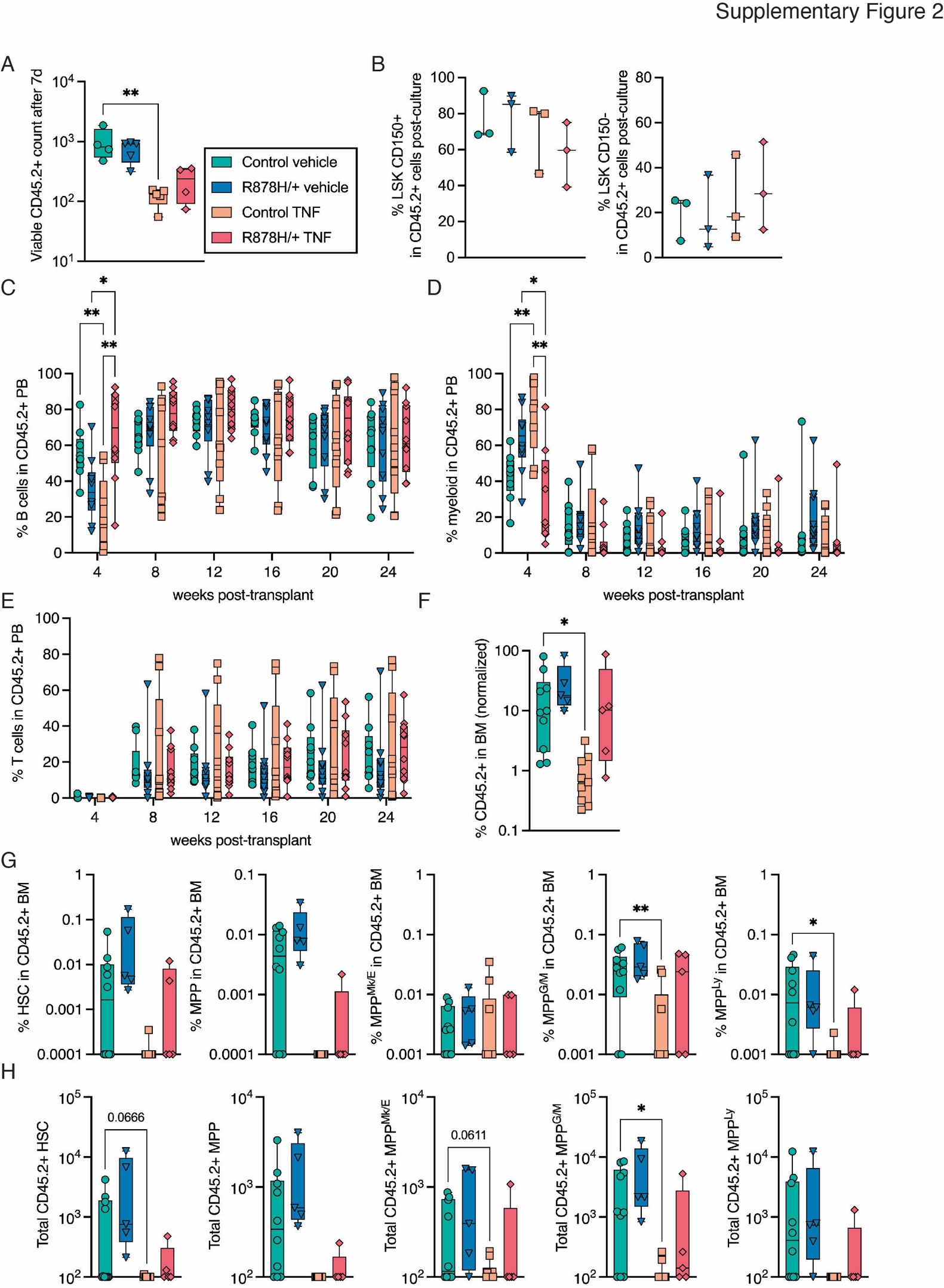

Supplementary Figure 2. Cell Type Composition of Control and *Dnmt3a*^R878H/+^ Hematopoiesis After *Ex Vivo* TNFα Stimulation. (A) Viable CD45.2+ cell count after 7 days of culture. Significance was calculated using one-way ANOVA with Tukey’s multiple comparisons test. (B) Frequency of cells with Lin- Sca+ Kit+ CD150+ (left) and Lin- Sca+ Kit+ CD150- (right) surface marker phenotypes after 7 days of culture. (C-E) Frequency of B cells (C), myeloid cells (D), and T cells (E) in donor derived PB. Significance was calculated using mixed-effects analysis with Tukey’s multiple comparisons test. (F) Frequency of donor cells in BM of recipient mice at 24-weeks post-transplant. Significance was calculated using Brown-Forsythe and Welch ANOVA with Welch’s correction. (G) Frequency and (H) total number of donor derived HSC, MPP, MPP^Mk/E^, MPP^G/M^, and MPP^Ly^ cells. Significance was calculated using Brown-Forsythe and Welch ANOVA with Welch’s correction. Dots represent individual recipient mice, boxes show 25 to 75^th^ percentile, line is median, whiskers show min to max. **P* < 0.05, ***P* < 0.01.

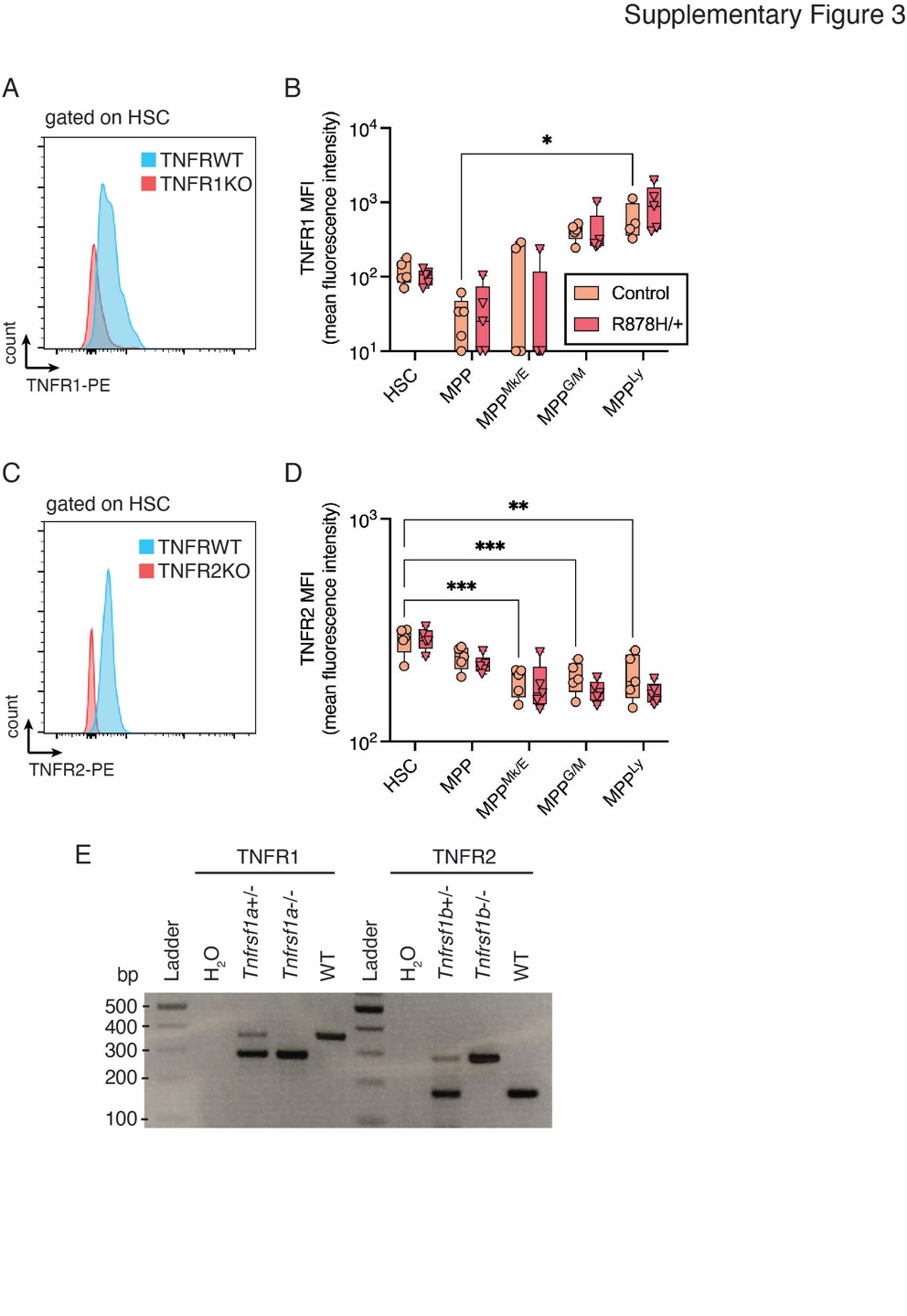

**Supplementary Figure 3. Flow Cytometric Analysis of TNFR1 and TNFR2 and TNFR Knockout Genotyping. (A)** Representative histogram of TNFR1 expression on HSCs compared to negative control (TNFR1 knockout HSCs). (**B**) MFI of TNFR1 on HSC, MPP, MPP^Mk/E^, MPP^G/M^, and MPP^Ly^ cells in Mx-Cre control vs. *Dnmt3a*^R878H/+^ (R878H/+) mice. Significance was calculated using two-way ANOVA with Tukey’s multiple comparisons test. (**C**) Representative histogram of TNFR2 expression on HSCs compared to negative control (TNFR2 knockout HSCs). (**D**) MFI of TNFR2 on HSC, MPP, MPP^Mk/E^, MPP^G/M^, and MPP^Ly^ cells in Mx-Cre control vs. *Dnmt3a*^R878H/+^ (R878H/+) mice. Significance was calculated using two-way ANOVA with Tukey’s multiple comparisons test. (**E**) PCR genotyping of BM cells isolated from *Tnfrsf1a* and *Tnfrsf1b* heterozygous and homozygous knockout animals. (**C, D**) Dots represent individual recipient mice, boxes show 25 to 75^th^ percentile, line is median, whiskers show min to max. **P* < 0.05, ***P* < 0.01, ****P* < 0.001.

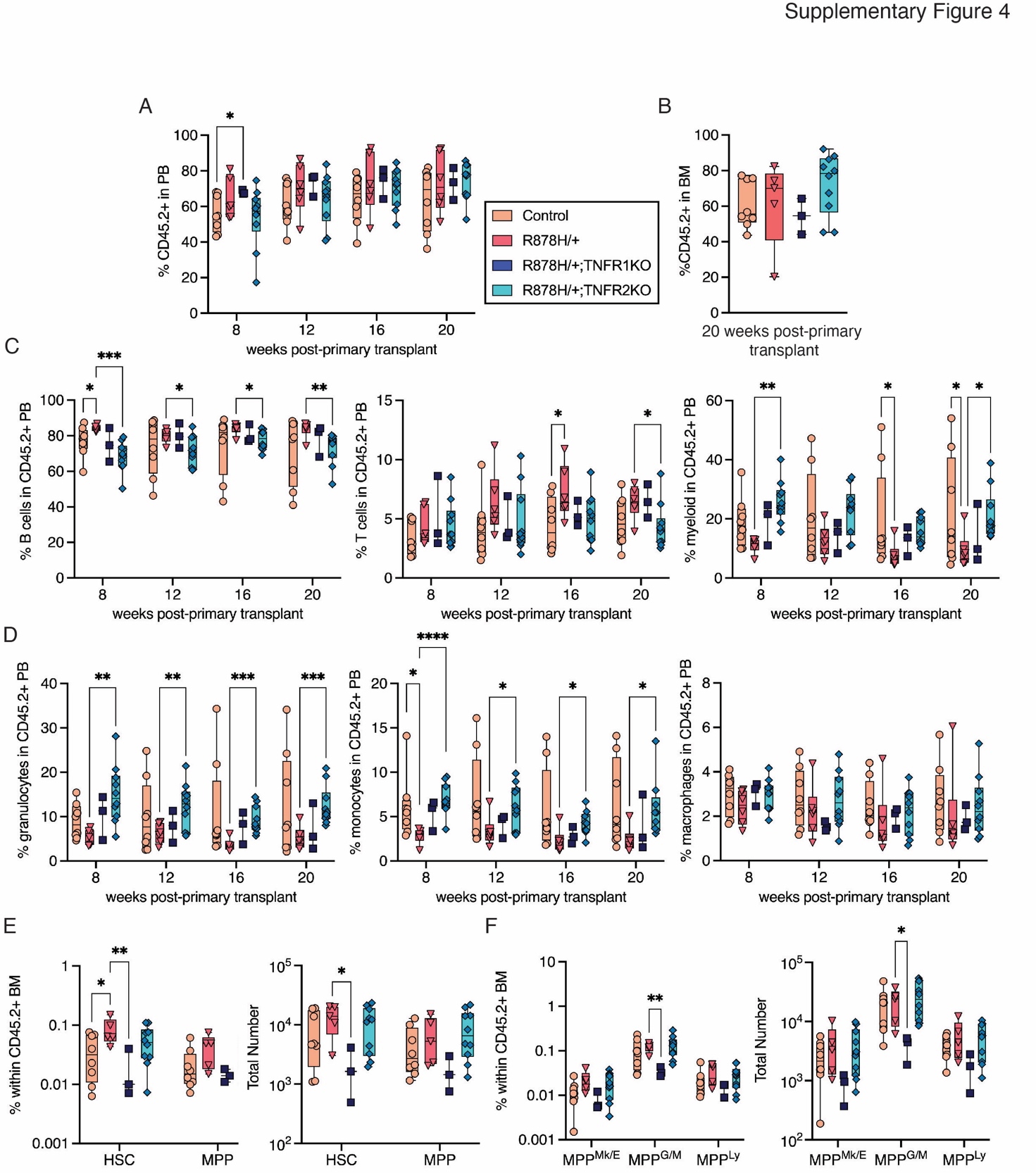

**Supplementary Figure 4. Multilineage Engraftment and Bone Marrow Analysis in Primary Transplant Recipient Mice.** (**A**) Frequency of donor cells in PB of primary transplant recipient mice. Significance was calculated using two-way ANOVA with Fisher’s LSD. (**B**) Frequency of donor cells in BM of primary transplant recipients at 20 weeks post-transplant. (**C**) Frequency of B cells (left), T cells (center) and myeloid cells (right) in donor derived PB. Significance was calculated using two-way ANOVA with Fisher’s LSD. (**D**) Frequency of granulocytes (left), monocytes (center) and macrophages (right) in donor derived PB. Significance was calculated using two-way ANOVA with Fisher’s LSD. (**E**) Frequency (left) and total number (right) of HSC and MPP cells in donor derived BM. Significance was calculated using two-way ANOVA with Fisher’s LSD. (**F**) Frequency (left) and total number (right) of MPP^Mk/E^, MPP^G/M^ and MPP^Ly^ cells in donor-derived BM. Significance was calculated using two-way ANOVA with Fisher’s LSD. Dots represent individual recipient mice, boxes show 25 to 75^th^ percentile, line is median, whiskers show min to max. **P* < 0.05, ***P* < 0.01, ****P* < 0.001, *****P* < 0.0001.

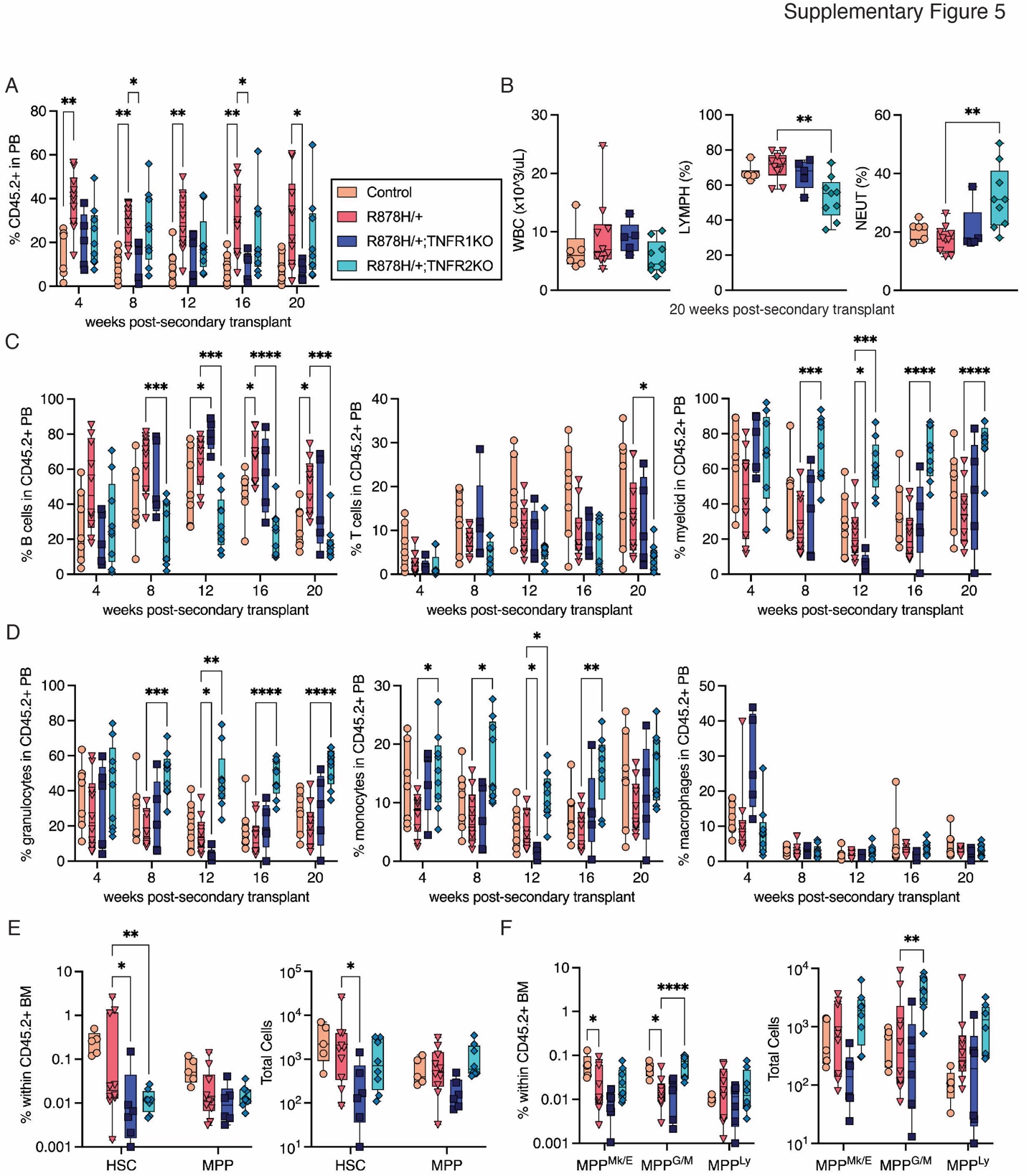

**Supplementary Figure 5.** **Multilineage Engraftment and Bone Marrow Analysis in Secondary Transplant Recipient Mice.** (**A**) Frequency of donor cells in PB of secondary transplant recipient mice. Significance was calculated using two-way ANOVA with Tukey’s multiple comparisons test. (**B**) White blood cell (WBC) counts and frequency of lymphocytes (LYMPH) and neutrophils (NEUT) in PB of secondary transplant mice at 20 weeks post-transplant. Significance was calculated using Brown Forsythe and Welch ANOVA with Welch’s correction. (**C**) Frequency of B cells (left), T cells (center) and myeloid cells (right) in donor derived PB. Significance was calculated using two-way ANOVA with Tukey’s multiple comparisons test. (**D**) Frequency of granulocytes (left), monocytes (center) and macrophages (right) in donor derived PB. Significance was calculated using two-way ANOVA with Tukey’s multiple comparison test. (**E**) Frequency (left) and total number (right) of HSC and MPP cells in donor derived BM. Significance was calculated using two-way ANOVA with Fisher’s LSD. (**F**) Frequency (left) and total number (right) of MPP^Mk/E^, MPP^G/M^ and MPP^Ly^ cells in donor-derived BM. Significance was calculated using two-way ANOVA with Fisher’s LSD. Dots represent individual recipient mice, boxes show 25 to 75^th^ percentile, line is median, whiskers show min to max. **P* < 0.05, ***P* < 0.01, ****P* < 0.001, *****P* < 0.0001.

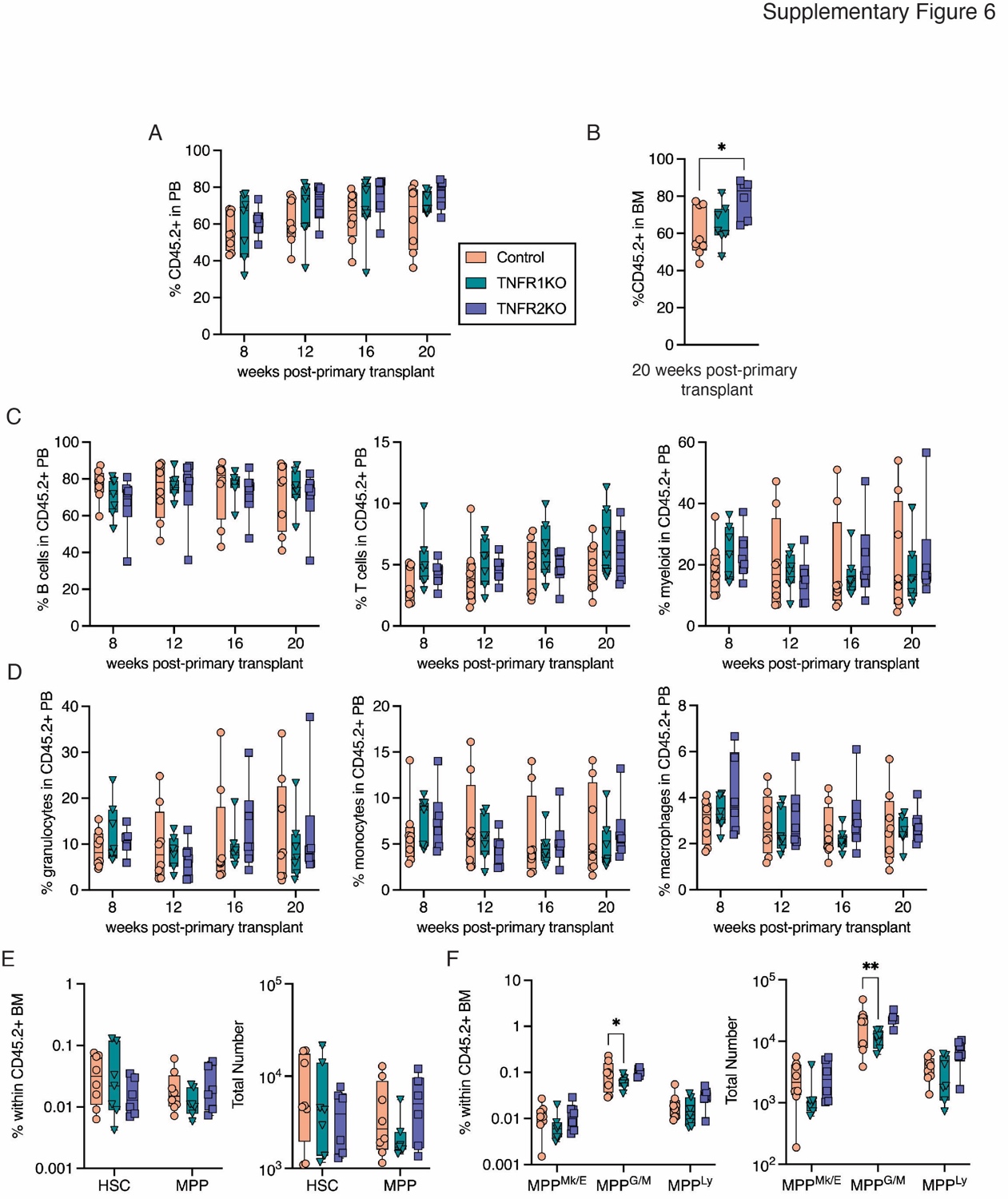

**Supplementary Figure 6. Multilineage Engraftment and Bone Marrow Analysis in Control Primary Transplant Recipient Mice.** (**A**) Frequency of donor cells in PB of primary transplant recipient mice. (**B**) Frequency of donor cells in BM of primary transplant recipients at 20 weeks post-transplant. Significance was calculated using one-way ANOVA with Holm-Sidak’s multiple comparisons test. (**C**) Frequency of B cells (left), T cells (center) and myeloid cells (right) in donor derived PB. (**D**) Frequency of granulocytes (left), monocytes (center) and macrophages (right) in donor d€ved PB. (**E**) Frequency (left) and total number (right) of HSC and MPP cells in donor derived BM. (**F**) Frequency (left) and total number (right) of MPP^Mk/E^, MPP^G/M^ and MPP^Ly^ cells in donor-derived BM. Significance was calculated using two-way ANOVA with Fisher’s LSD. Dots represent individual recipient mice, boxes show 25 to 75^th^ percentile, line is median, whiskers show min to max. **P* < 0.05, ***P* < 0.01.

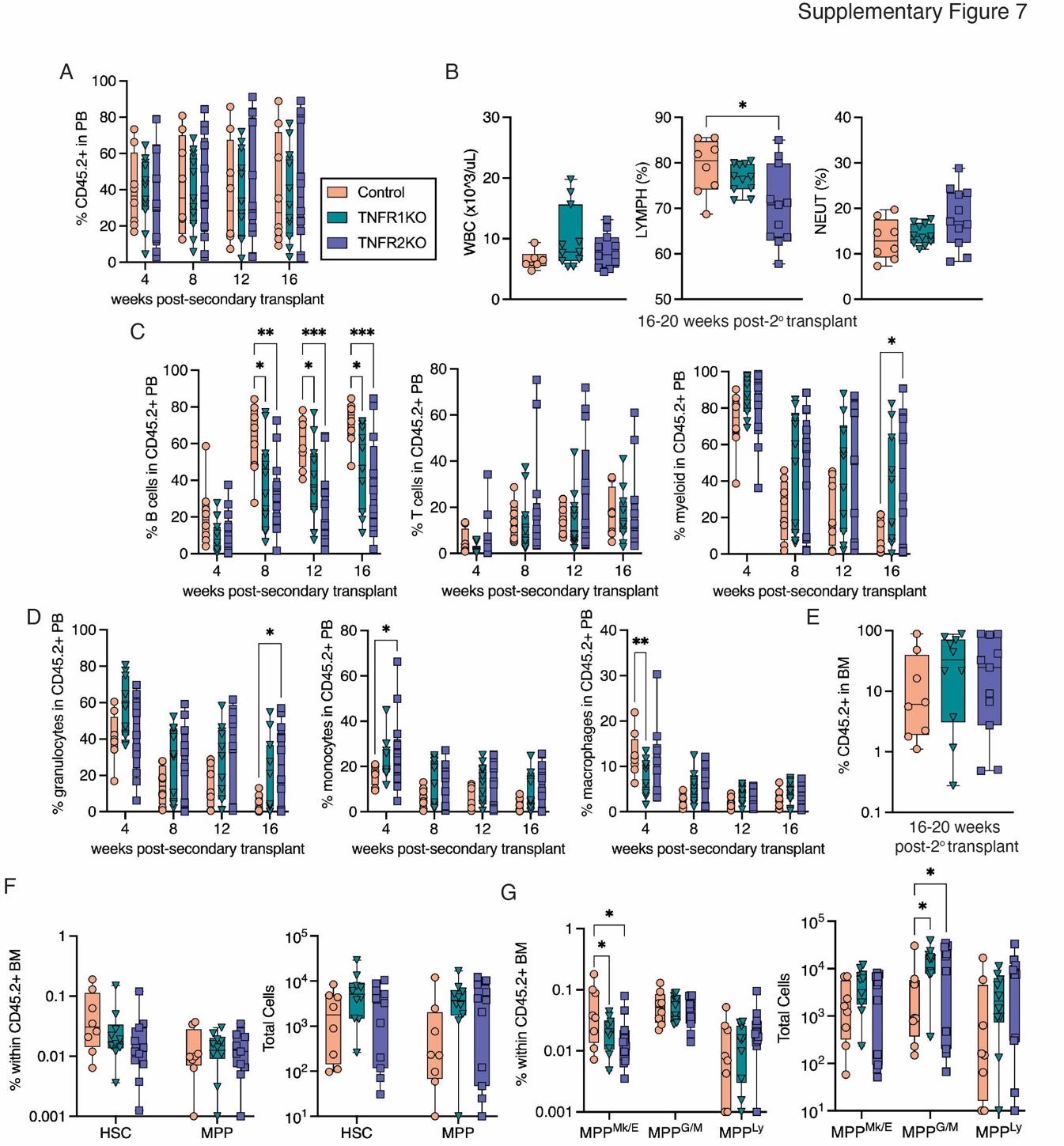

**Supplementary Figure 7.** **Multilineage Engraftment and Bone Marrow Analysis in Control Secondary Transplant Recipient Mice.** (**A**) Frequency of donor cells in PB of secondary transplant recipient mice. (**B**) White blood cell (WBC) counts and frequency of lymphocytes (LYMPH) and neutrophils (NEUT) in PB of secondary transplant mice at 16-20 weeks post-transplant. Significance was calculated using Brown Forsythe and Welch ANOVA with Welch’s correction. (**C**) Frequency of B cells (left), T cells (center) and myeloid cells (right) in donor derived PB. Significance was calculated using two-way ANOVA with Tukey’s multiple comparisons test. (**D**) Frequency of granulocytes (left), monocytes (center) and macrophages (right) in donor derived PB. Significance was calculated using two-way ANOVA with Tukey’s multiple comparisons test. (**E**) Frequency of donor cells in BM of primary transplant recipients at 16-20 weeks post-transplant. (**F**) Frequency (left) and total number (right) of HSC and MPP cells in donor derived BM. (**G**) Frequency (left) and total number (right) of MPP^Mk/E^, MPP^G/M^ and MPP^Ly^ cells in donor-derived BM. Significance was calculated using two-way ANOVA with Tukey’s multiple comparisons test. Dots represent individual recipient mice, boxes show 25 to 75^th^ percentile, line is median, whiskers show min to max. **P* < 0.05, ***P* < 0.01, ****P* < 0.001.

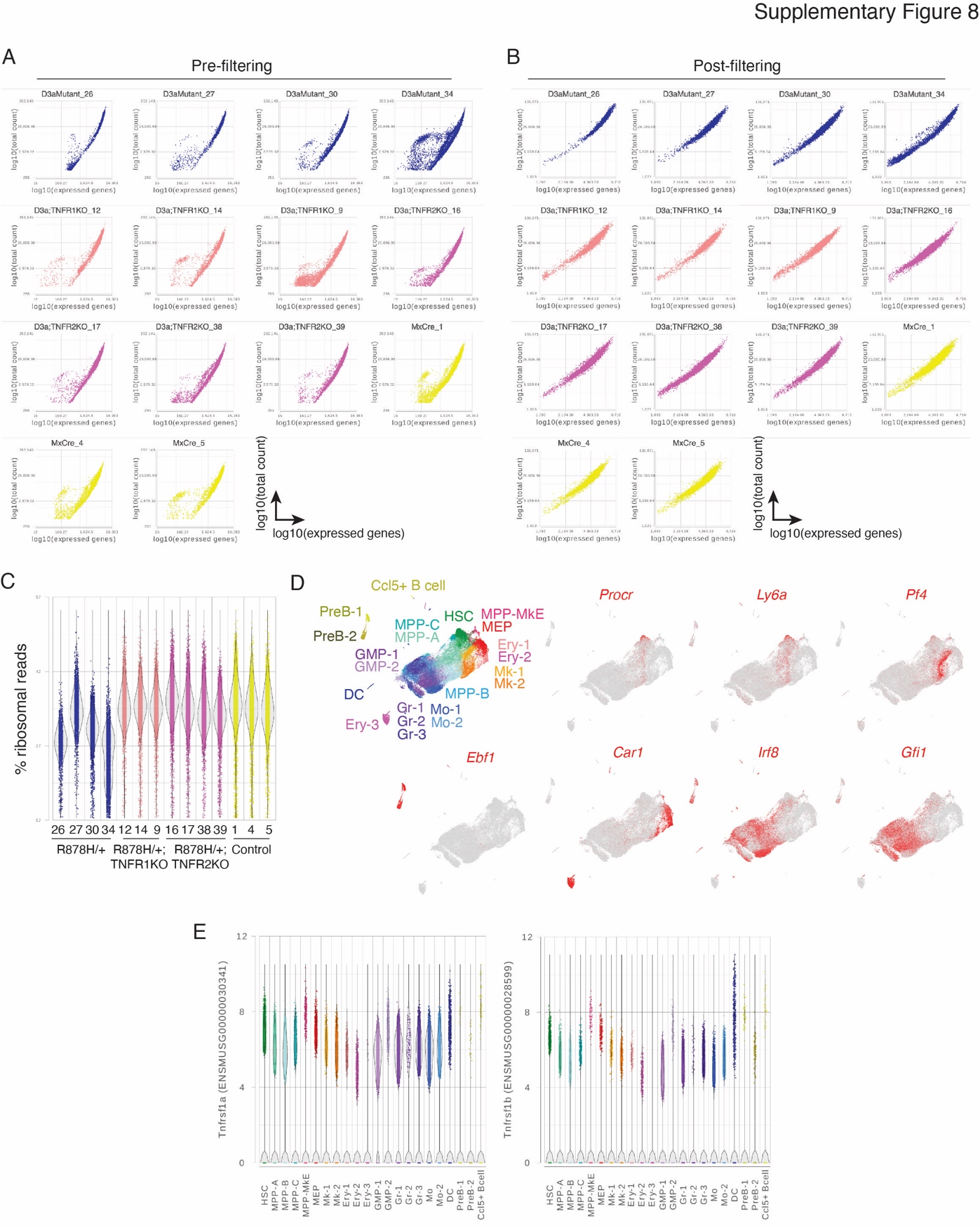

**Supplementary Figure 8. Quality Control and Clustering of Single Cell RNA-seq Data.** (**A, B**) Individual biological replicate single cell RNA-seq samples prior to (**A**) and following (**B**) filtering based on number of expressed genes, mitochondrial, and ribosomal reads. (**C)** Frequency of ribosomal reads in each library after filtering. (**D**) UMAP projection of combined data identifying 22 cell clusters and showing representative marker gene expression defining cell types including *Procr* and *Ly6a* (HSC), *Pf4* (megakaryocyte/Mk), *Ebf1* (B cells/preB), *Car1* (erythroid/Ery), *Irf8* (monocyte/Mo), and *Gfi1* (granulocyte/Gr). (**E**) Expression of *Tnfrsf1a* and *Tnfrsf1b* in individual cells across 22 cell clusters.

**Supplementary Table 1. Marker Genes Used to Classify 22 Cell Clusters Identified by Single Cell RNA-seq Analysis** (see Excel file).
